# Pleiotropic effects of a recessive C*OL1α2* mutation occurring in a mouse model of severe osteogenesis imperfecta

**DOI:** 10.1101/2024.02.15.580510

**Authors:** Michelangelo Corcelli, Rachel Sagar, Ellen Petzendorfer, Mohammad Mehedi Hasan, Hilda I de Vries, Fleur S van Dijk, Anna L David, Pascale V Guillot

## Abstract

Approximately 85-90% of individuals with Osteogenesis Imperfecta (OI) have dominant pathogenic variants in the *COL1A1 or COL1A2* genes. This leads to decreased or abnormal Collagen type I production. Subsequently, bone formation is strongly reduced, causing bone fragility and liability to fractures throughout life. OI is clinically classified in 5 types with the severity ranging from mild to lethal depending on the gene and the type and location of the OI-causative variant and the subsequent effect on (pro) collagen type I synthesis. However, the specific effects on the phenotype and function of osteoblasts are not fully understood.

To investigate this, the OI murine model was used, with the *oim*/*oim* (OIM) mice closest resembling severely deforming OI type 3 in humans. We showed that in OIM, COL1 mutation results in a multifactorial inhibition of the osteogenic differentiation and maturation as well as inhibition of osteoclastogenesis. The phenotype of differentiated OIM osteoblasts also differs from that of wild type mature osteoblasts, with upregulated oxidative cell stress and autophagy pathways, possibly in response to the intracellular accumulation of type I collagen mRNA. The extracellular accumulation of defective type I collagen fibres contributes to activation of the TGF-β signalling pathway and activates the inflammatory pathway. These effects combine to destabilise the balance of bone turnover, increasing bone fragility. Together, these findings identify the complex mechanisms underlying OI bone fragility in the OIM model of severe OI and can potentially enable identification of clinically relevant endpoints to assess the efficacy of innovative pro-osteogenic treatment for patients with OI.

---

Bones provide structural support to the body, protection of vital organs, function as the main site of hematopoiesis, and play a key role in the regulation and storage/release of minerals, growth factors and hormones.^1^

Osteogenesis imperfecta (OI), or brittle bone disease, is a rare monogenic inherited connective tissue disorder with a prevalence of 6-7/100,000. The main feature of OI is increased liability to fractures throughout life, osteopenia, skeletal deformity, and other features including blue sclerae, hearing loss, dentinogenesis imperfecta and short stature.^2^ OI is clinically classified into five types (OI type 1-5). Approximately 85-90% of individuals with OI have a heterozygous pathogenic variant in the *COL1A1* or *COL1A2* gene encoding respectively type I (pro)collagen alpha 1 or alpha 2 chain. These chains intertwine in a triple helix structure to form (pro)collagen type I. Depending on the gene involved, the location and the type of variant, there is either a 50% decrease of (pro)collagen type I synthesis which leads to OI type 1 (haploinsufficiency) or there is production of abnormal (pro)collagen type I which usually leads to OI types 2-4. The most common types of variants causing *COL1A1*/2 related OI type 3, are glycine substitutions as (pro)collagen type I has a large, conserved domain consisting of Gly-X-Y triplets. Glycine is the smallest amino acid and when substituted for a different amino acid this hampers proper folding and assembly of the triple helix^3,4^ which results in OI types 2-4. Currently, more than 1000 different pathogenic variants have been reported in *COL1A1*/*COL1A2*.^2^

In OIM (osteogenesis imperfecta murine) mice (*B6C3fe-a/a-oim)*, a recessive point mutation (a G deletion at nucleotide 3983 in *COL1A2*) in the gene coding for the alpha 2 chain of type I (pro)collagen (*COL1A2*) results in an alteration of the sequence of the last 48 amino acids, leading to the absence of COL1A2 protein, despite mRNA transcription. In homozygous OIM (*oim/oim*), this leads to the formation of homotrimeric α1(I)_3_ type I collagen, instead of the normal heterotrimeric collagen α1(I)_2_α2(I)_1_. This recessive variant has the same effect in humans, namely structurally abnormal collagen type I being produced. The impairment of OIM osteoblastic differentiation and the increased osteoclast formation^5,6^ as well as the activation of the TGF-β signalling pathway^7^ have previously been reported.^8^ Using the α2(I)-G610C mouse model of OI, Mirigian et al.^9^ also showed that osteoblast malfunction appeared to be caused by the cell stress response to procollagen misfolding.

To propose a theoretical model of OI bone fragility, we used primary pre-osteoblasts isolated from the calvaria of homozygous OIM and wild type (WT) neonatal mice to determine the genetic impact of the recessive variant on osteoblast phenotype during the process of osteogenic differentiation.

## Results

### Micro computed tomography (microCT) of neonate OIM and WT proximal tibia

Qualitatively, comparative analysis of WT and OIM proximal tibia in 1-week-old mice (**Figure 1**) and in 8-week-old mice (Figure 2) by microCT clearly shows the taller metaphysis in younger mice, along with a smaller medullary canal thickness. In adult OIM mice, we observed the typical decrease in cortical thickness and trabecular network, which are the hallmark of the OIM phenotype.

**Figure 1.**
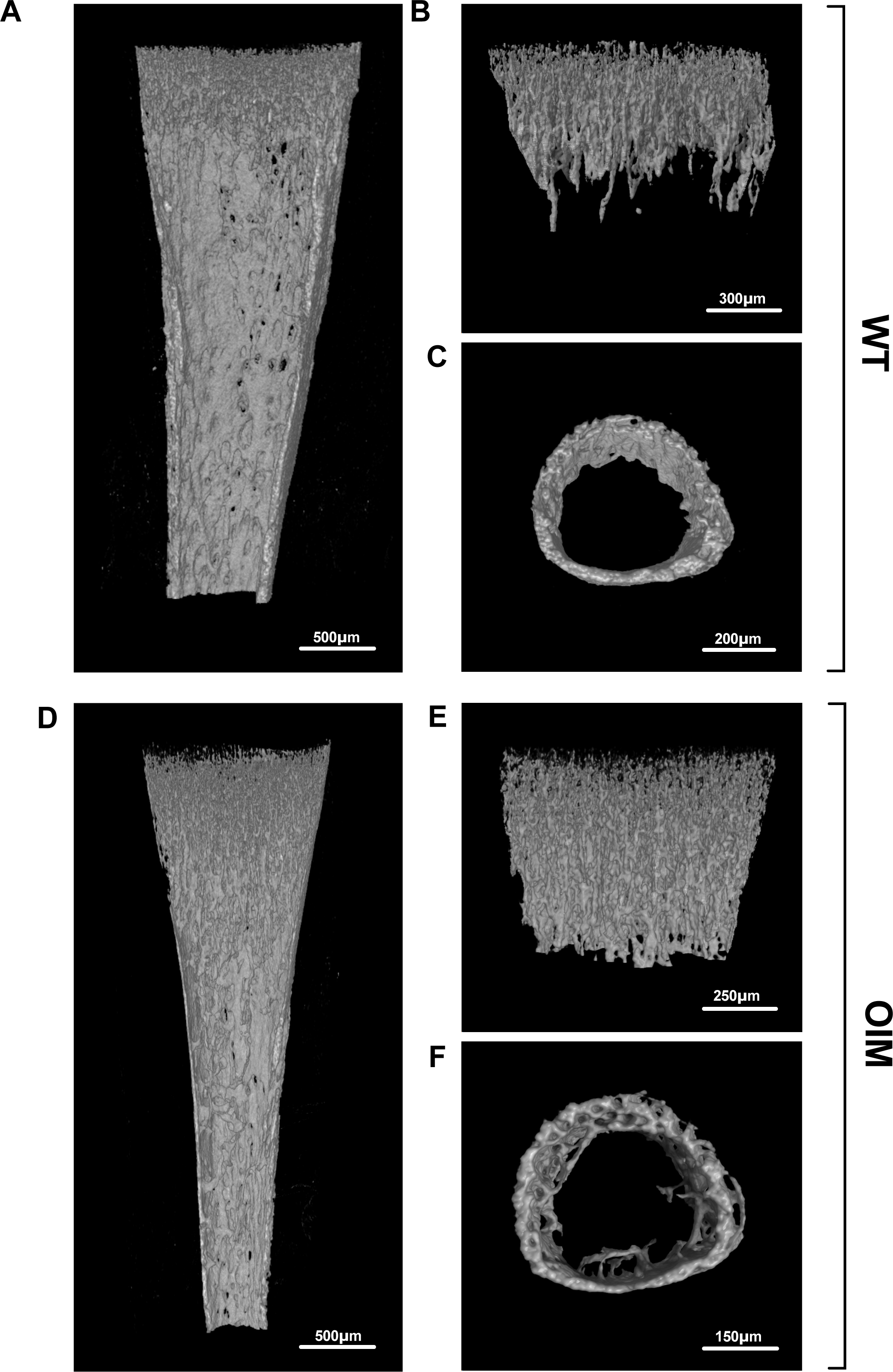
X-ray microCT imaging of the proximal tibia from 7 days old mice. **(A, D)** Representative sagittal sections of the proximal tibia showing the trabecular and cortical bone architecture. **(B, E)** Segmented view of the trabecular bone at proximal metaphysis obtained from 7-day old mice (WT, top and OIM, bottom). **(C, F)** Representative microCT image of the cortical bone at mid-diaphysis.

**Figure 2.**
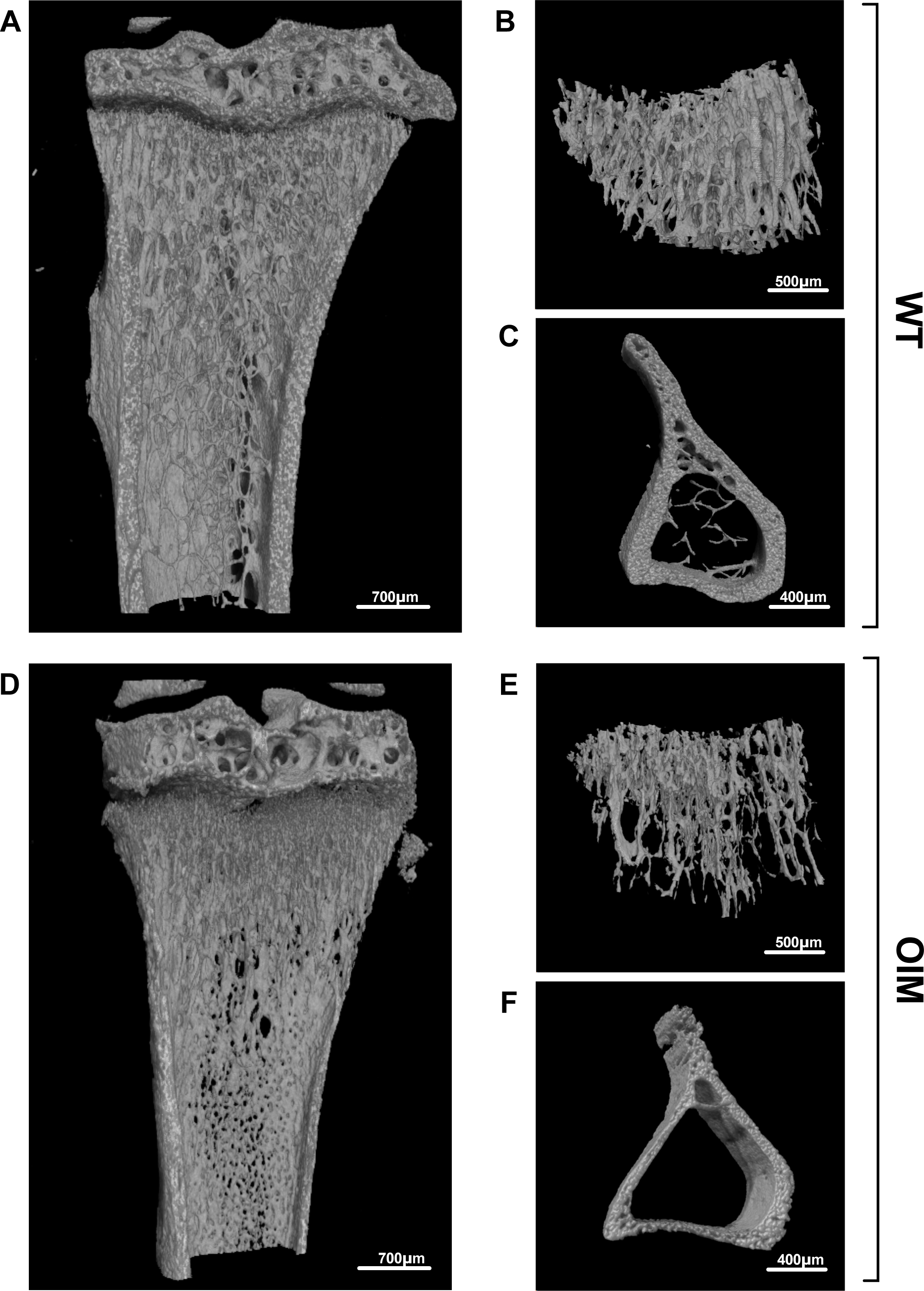
X-ray microCT imaging of the proximal tibia from 8 weeks-old mice. **(A, D)** Representative sagittal sections of the proximal tibia showing the trabecular and cortical bone architecture. **(B, E)** Segmented view of the trabecular bone obtained from 8-week-old mice (WT, top OIM, bottom). **(C, F)** Representative microCT image of the cortical bone at mid-diaphysis obtained from 7-day old mice.

Quantitatively, compared to their WT counterparts, proximal tibia analysis of 1-week-old OIM revealed a significant decrease in medullary (or marrow) canal thickness (which measures the diameter of the medullary canal) i.e., 0.40±0.01 mm vs 0.33±0.02 mm, WT vs OIM, mean±SEM, n=7, P<0.1), a significant increase in TMD (tissue mineral density) i.e., 0.89±0.01 g/cm^2^ vs 0.80±0.10 g/cm^2^, n=7, P<0.0001), and a significant decrease in medullary volume i.e., 0.15±0.01 mm^3^ vs 0.1±0.01 mm^3^, n=7, P,0.1) with no significant differences for cortical volume, cortical thickness and total porosity (**Figure 1** and **Figure 3A**). Comparative analysis of trabecular bone (**Figure 1** and **Figure 3B**) showed no significant difference for trabecular thickness, intersection surface (which is the surface of the volume of interest intersected by solid binarized objects) and trabecular pattern factor (which relates to the arrangement and connectivity of trabeculae within bone). However, OIM bones showed a significant increase in BV/TV (which corresponds to the bone volume fraction, calculated by the ratio of the segmented bone volume to the total volume of the region of interest) i.e., 7.52±0.5 % vs 11.08±0.37 %, n=7, P<0.0001), in BS/TV (which quantifies the bone surface density, as the ratio of the segmented bone surface to the total volume of the region of interest) i.e., 21.13±1.16 mm^-1^ vs 30.96±0.86 mm^-1^, n=7, P<0.0001) and in trabecular number (5.57±0.32 mm^-1^ vs 8.19±0.23 mm^-1^, n=7, P<0.0001).

**Figure 3.**
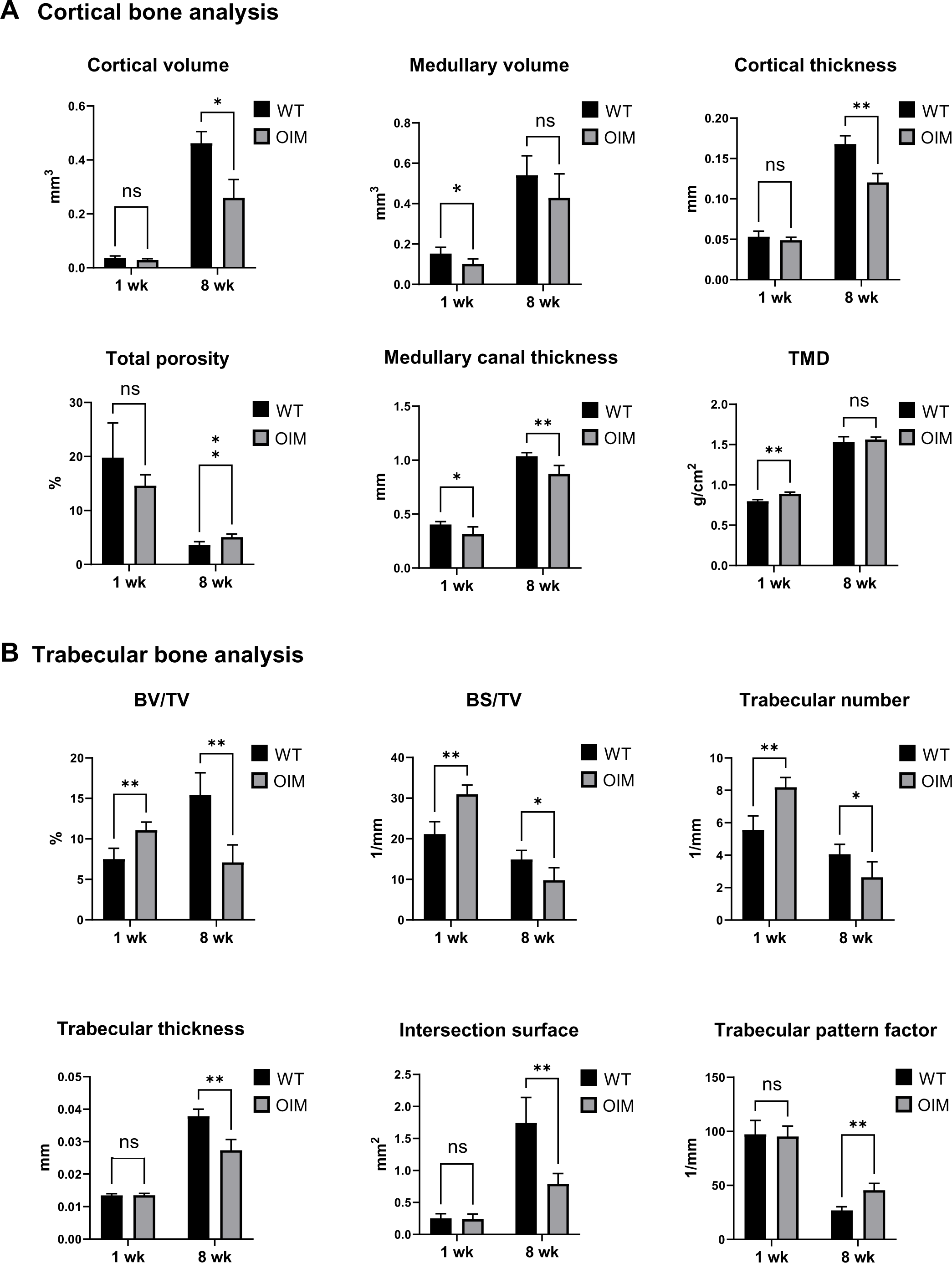
Micro CT morphometric analysis. **(A)** Quantification of morphometric parameters of tibial trabecular bone in 1 week-old and in 8 week-old WT and OIM mice. Cortical volume (mm^3^), medullary volume (mm^3^), cortical thickness (mm), total porosity (%), medullary canal thickness (mm), tissue mineral density (TMD, g/cm^2^). **(B)** Quantification of morphometric parameters of tibial cortical bone. Bone volume percent (BV/TV, %), bone surface density (BS/TV, 1/mm), trabecular number (1/mm), trabecular thickness (mm), Intersection surface (mm^2^), trabecular pattern factor (1/mm). All microCT parameters were analysed using Bonferroni’s multiple comparison post hoc test., ***P□<□0.001, **P□<□0.01 and * P□<□0.05. ns: not significant. n=7 for all the groups.

In contrast, and as expected, in 8-week-old mice, we observed a decrease in cortical volume in OIM mice compared to WT (0.43±0.02 mm^3^ vs 0.26±0.03 mm^3^, n=7, P<0.0001), cortical thickness (0.17±0.01 mm vs 0.12±0.01 mm, n=7, P<0.0001), medullary canal thickness (1.04±0.01 mm vs 0.88±0.03 mm, n=7, P<0.001), BV/TV (15.40±1.13 vs 7.10±0.88, n=7, P<0.001), BS/TV (14.90±0.90 1/mm vs 9.81±1.25 1/mm, n=7, P<0.01), trabecular number (4.06±0.25 1/mm vs 2.64±0.39 1/mm, n=7, P<0.01), trabecular thickness (0.04±0.00 mm vs 0.03±0.00 mm, n=7, P<0.0001), intersection surface (1.75±0.16 mm^2^ vs 0.79±0.06 mm^2^, n=7, P<0.001), and an increase in total bone porosity (3.61±0.25 % vs 5.10±0.23 %, n=7, P<0.001). In addition, as we previously reported, we observed an increase in trabecular pattern factor (26.92±1.40 mm^-1^ vs 45.62±2.52 mm^-1^, n=7, P<0.0001), indicative of a less interconnected and structurally sound trabecular network, which is associated with lower bone strength and compromised bone integrity.

### *In vitro* mineralization deposition by OIM osteoblasts is abundant but disorganized

Primary pre-osteoblasts isolated from the calvaria of WT and OIM neonates, were expanded in non-osteogenic media (**Figure 4A**). Upon reaching near confluence, their expansion medium was replaced by osteogenic inductive medium and a 30-day differentiation period commenced, during which medium was replaced three times per week (**Figure 4B**). Both cell types adopted the typical cobblestone morphology, which was qualitatively visible from as early as day 21, whereby WT cells deposited minerals in a very structured manner, forming spicules with well-defined thick borders (**Figure 4B**). By day 30, these spicules became interconnected to form a network of well-organised mineral matrix. In contrast, OIM cells produced an abundant amount of minerals that failed to organize. Instead, they remained diffuse throughout the dish and did not form clear mineralization spicules with well-defined borders. Alizarin red staining, which detects the presence of calcium deposits in a mineralized matrix, confirmed the formation of mineralized spicules by WT osteoblasts and the decreased organized deposition of calcium by OIM cells (**Figure 4C**), despite the total amount of minerals not significantly differing (**Figure 4D**).

**Figure 4.**
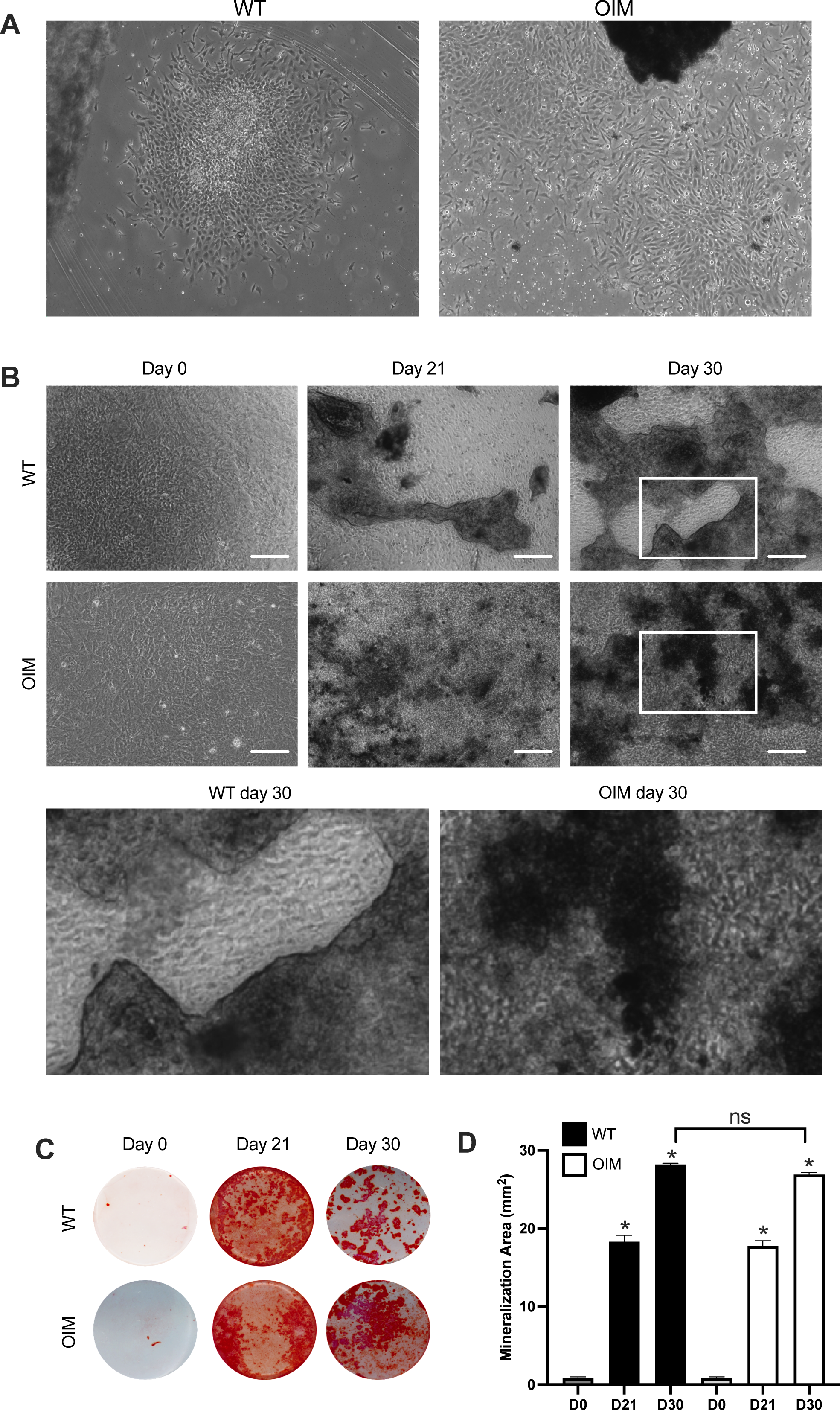
Invitro assessment of the mineral deposition in WT and OIM osteoblasts. **(A)** Phase contrast of primary calvaria osteoblasts. **(B)** Phase contrast of osteoblasts cultured in basal and osteogenic conditions at day 0, 21, and 30. **(C)** Alizarin Red staining. **(D)** Measurement of the mineralization area.

### Comparative analysis of OIM and WT pre-osteoblast differentiation down the osteogenic pathway showed that the OIM-causative mutation triggers cell stress and inflammation

Whole genome transcriptome comparative analysis (RNA-Seq) of WT and OIM cells after 0 or 21 days of culture in 2D osteogenic permissive conditions provides insight as to whether the recessive OIM-causative variant contributes, directly or indirectly, to the transcriptional regulation of the osteogenic and metabolic pathways (**Figure 5**). Here, we are comparing how both cell types (WT and OIM) respond to osteogenic induction over a 21-day period. In OIM cells, the recessive *CoL1a2* variant affects the reading frame and resulted in an incorrect pro-peptide and a new stop codon.^10^ This lead to the intracellular accumulation of a defective a1(I)_2_ mRNA transcript (non-sense mediated decay escape). In the bone extracellular matrix (ECM), this results in bone tissue accumulation of a dysfunctional type I collagen homotrimer a1(I)_3_, in replacement of the wild type a1(I)_2_a2(I)_1_ heterotrimer.^11^ Our data reveals the upregulation of genes involved in the production of reactive oxygen species (ROS), such as *XDH* (xanthine dehydrogenase), which causes oxidative stress and inflammation in response to defective mRNA and/or the accumulation of dysfunctional proteins.^12^ Cellular inflammation and cell stress in OIM cells is demonstrated by the upregulation of *NOS2* (nitric oxide synthase 2), which is known to be induced in response to inflammatory signals as well as *DAPK1* (death associated protein kinase 1) and *TNFSF10* (tumour necrosis factor related apoptosis inducing ligand), acting as a pro-apoptotic and pro-autophagic protein during cellular stress.^13^

**Figure 5.**
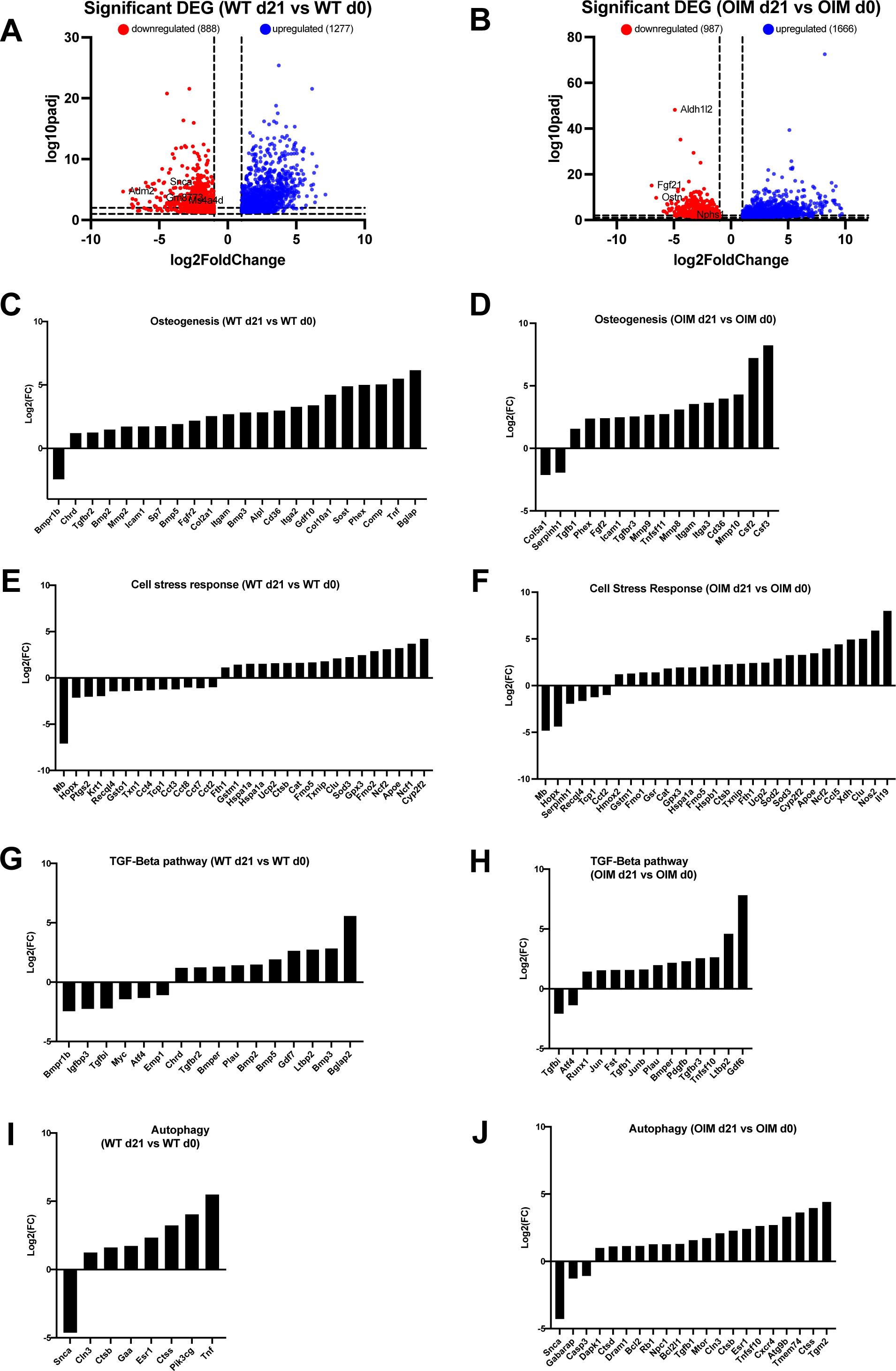
RNAseq analysis of WT mature osteoblasts vs WT pre-osteoblasts and OIM mature osteoblasts vs OIM pre-osteoblasts during osteogenic differentiation. (A) Volcano plots showing significant (FDR adjusted p<0.05) DEGs (differentially expressed genes) in mature WT (WT d21) vs WT pr-osteoblasts (WT d0) (A) and mature OIM (OIM d21) vs OIM pr-osteoblasts (OIM d0) (B) after 3 weeks in osteogenic induction medium. DEGs were plotted by pathways: osteogenesis (C, D), cell stress response (E, F), TGF-Beta pathway (G, H), autophagy (I, J).

### Cell stress in OIM cells activates pro-apoptotic and pro-autophagic pathways

The activation of the pro-apoptotic and pro-autophagic pathways in OIM cells acts as a protective mechanism against oxidative stress and other cellular stressors such as the intracellular presence of defective mRNA. For example, in OIM cells, we observed the upregulation of *RB1* (retinoblastoma 1), which plays a role in regulating apoptosis.^14^ Activation of the pro-autophagy pathway in OIM is evidenced by the upregulation of *DRAM1* (damage-regulated autophagy modulator 1), *ATG9b* (autophagy related protein 9b) and *TGM2* (trans glutaminase 2), the expression of which is induced by oxidative stress and which contributes to the initiation of autophagy by promoting the formation and expansion of the autophagosome^15–17^. *CXCR4* (C-X-C chemokine receptor type 4), which is part of the cellular stress response and which expression has been shown to increase autophagy activity^18^, is also upregulated. Activation of autophagy in OIM cells is also evidenced by the upregulation of *TMEM74* (transmembrane 74), which promotes autophagy.^19^

### Antioxidant defence mechanism in OIM cells

However, OIM cells may simultaneously be protected from oxidative damage, as evidenced for example by the upregulation of *GSR* (glutathione reductase), which plays a vital role in maintaining the cellular antioxidant defence mechanisms^20^, and *HMOX2* (heme oxygenase 2), which protects cells against oxidative stress and inflammation through its involvement in heme degradation and subsequent bilirubin-mediated neutralization of ROS.^21^ OIM cell protection against ROS-induced intra-cellular damage is also suggested by the upregulation of *SOD2* (superoxide dismutase 2), a mitochondrial enzyme that maintains the cellular redox balance by scavenging superoxide radicals and thereby preventing apoptosis triggered by excessive ROS.^22^

Promotion of OIM cell survival as a defence mechanism against oxidative stress-induced apoptosis is also evidenced by the upregulation of genes from the Bcl-2 family, which are involved in the regulation of apoptosis. For example, we observed an upregulation of *BCLl2* (B cell lymphoma 2), and *BCL2L1* (B-cell lymphoma 2-like 1), which promote cell survival by preventing the degradation of various cellular components, and by decreasing autophagy through the inhibition of *BECN1* (beclin-1), concomitant with the downregulation of *CASP3* (caspase 3), a key mediator of apoptosis.^23^ Another illustration of the cellular defence mechanism activated in OIM cells to counteract oxidative stress-induced apoptosis and autophagy is the upregulation of *mTOR* (mammalian target of rapamycin), which leads to the suppression of key autophagy-related proteins that are essential for initiating the autophagic process.^24^

### Activation of the TGF-β pathway in OIM stimulates the pathway involved in the proliferation of osteoprogenitors and modulation of bone resorption

Excessive TGF-β (transforming growth factor beta) signalling is the most significant pathogenic mechanisms in OI.^25^ The abnormal bone ECM structure prevents the proper decorin-mediated sequestration of latent TFG-β in the bone matrix, which results in excessive release of TFG-β in its active form.^7^ The upregulation of the TFG-β pathway in OIM cells was evidenced by the increase in the expression of *TGF-β 1* and *TGF-β -R3* (transforming growth factor beta receptor 3) which are both involved in the recruitment of endogenous MSCs and the proliferation of osteoprogenitor cells.^26^ TGF-β3 promotes osteoprogenitor cell proliferation through the activation of non-Smad pathways such as MAPK and P13K, which stimulate the production of growth factors, but is also secreted by osteoprogenitors themselves in an autocrine manner. Interestingly, *MAPK13* is upregulated in OIM cells but Smad-related genes are not, suggesting that *TGFb-R3* modulates osteogenesis through the MAPK pathway. Finally, *TFG-β-R1*, which was upregulated in OIM, is directly responsible for initiating the intracellular signalling cascade upon binding of TFG-β ligands, whilst *TFG-β-R2*, which is upregulated in WT but not in OIM, forms a complex with *TFG-β-R1* upon ligand binding. The upregulation of *PDGFB* (platelet derived growth factor B) and *JUNB* (JunB proto-oncogene or AP-1 transcription factor subunit) further suggest the proliferation of OIM osteoprogenitors.^27,28^

### Activation of the TGF-β pathway in OIM impacts osteogenesis by inhibiting both osteoblast initiation and maturation

We observed that *GDF6* (Growth differentiation factor 6, also called BMP13, bone morphogenic protein 13) is strongly upregulated in OIM whilst *GDF7* (BMP12) is upregulated in WT. Interestingly, GDF6 is a potent inhibitor of bone formation and inhibits the expression of late markers of osteogenic differentiation.^29^ Conversely, GDF7 induces the expression of osteogenic key genes involved in the mineralization of the bone matrix^30^ including *ALPL* (alkaline phosphatase), which facilitates the formation of hydroxyapatite crystals, and *BGLAP* (osteocalcin), which stimulates calcium ion deposition and matrix hardening. Additional genes involved in osteoblast maturation which are expressed by WT but not OIM cells include *SOST* (Sclerostin), *FGFR2* (Fibroblast Growth Factor Receptor 2) and *SP7* (Osterix). SOST is expressed by osteocytes and regulates the balance between bone formation and bone resorption by inhibiting early osteoblast activity.^31^ Osterix is a transcription factor that plays a role in the maturation of pre-osteoblasts into mature osteoblasts by activating *BGLAP* and *ALPL*, as well as in bone mineralization through the deposition of calcium and phosphate ions into the collagen matrix. The fact that *SP7* is upregulated in WT but not in OIM provides additional evidence that OIM have compromised osteoblast maturation and matrix mineralization. In addition to promoting expression of *SP7*, FGFR2 promotes osteoblast differentiation, indicating that the initiation of osteoblast differentiation may also be compromised in OIM.^32^

### Collagen’s functionality and matrix mineralization are compromised in OIM

*SERPINH1* (collagen binding protein HSP47), which plays a crucial role in the proper folding and stabilization of collagen molecules by binding to newly synthesized collagen chains in the endoplasmic reticulum (ER)^33^, is downregulated in OIM cells, suggesting that homotrimeric type I collagen’s stability and functionality may be compromised in the ECM. Although *PHEX* (phosphate regulating neutral endo peptidase) is upregulated in both WT and OIM, upregulation is markedly greater in WT, further evidencing the impairment of bone mineralization in OIM, since PHEX regulates phosphate levels in the body, which is essential for the formation of hydroxyapatites.^34^ *ITGA 2 and 3* genes (integrin alpha-2 or 3) have different functions. *Itga-2*, which is upregulated in WT only, is involved in the bonding of osteoblasts to collagen fibres in the bone ECM. This process ensures the proper function of osteoblasts.^35^ On the contrary, *Itga-3*, which is upregulated in OIM only, is involved in the adhesion of osteoblasts to non-collagenous ECM proteins.^35^

### Modulation of the BMP pathway in OIM contributes to the production of a defective bone ECM, impedes the proper transition to mature osteoblasts and impairs bone remodelling

*BMPR1B* (bone morphogenic protein receptor 1b)’s downregulation in WT cells, but not in OIM, ensures the proper commitment to osteogenesis and is involved in the transition to a mature osteoblast stage, as mature osteoblast do not depend on BMP signalling in the same way as progenitor cells. *BMPR1B* is also involved in a negative feedback loop to fine-tune BMP signalling. In line with these results, we observed an upregulation of *CHRD* (chordin) in WT only. Chordin regulates the activity of BMPs and acts as a BMP antagonist by binding to BMPR1B, thereby ensuring that BMPs are not excessively activated during the early stages of osteogenesis.^36^ *GDF10* (growth differentiation factor 10), which also acts as an antagonist to certain BMPs, is only upregulated in WT, further suggesting that bone remodelling is impaired in OIM. We observed that *BMP-2,3,5* genes^37^ are upregulated in WT but not OIM. BMP-2 promotes the expression of *SP7*, which is only upregulated in WT cells, and both enhances the production of bone ECM and plays a crucial role in matrix mineralization.

### Osteoclastogenesis and bone ECM degradation are stimulated in OIM

*TNF* (Tumour Necrosis Factor) is upregulated in WT only, further suggesting that bone remodelling is unbalanced in OIM, as TNF influences osteogenesis by regulating bone turnover and stimulating osteoblasts to produce factors such as VEGF (Vascular endothelial growth factor). *MMP2,8,9,10* (Matrix Metallo Proteinases) are responsible for degradation of various components of the extracellular matrix. More precisely, MMP2, which is upregulated in WT but not in OIM, is involved in the degradation of ECM components, breaking down collagen to allow osteoclasts to remove old bone tissue, indicating that OIM may fail to degrade dysfunctional collagen molecules.^38^ However, MMP8 and MMP10 which are also involved on collagen degradation in response to inflammation,^39^ are only upregulated in OIM. Similarly, MMP9, which is upregulated in OIM, is involved in bone resorption by helping osteoclasts assemble in the resorption pit.^40^ *TNFSF11* (tumour necrosis factor ligand superfamily 11, also called RANKL or receptor activator of nuclear factor kappa-B ligand), which is a key regulator of osteoclast differentiation and bone resorption, is only upregulated in OIM.^41^ Other genes stimulating osteoclastogenesis under the influence of *TNFSF11*, are *CSF2* and *CSF3* (colony stimulating factors 2 and 3), which are only upregulated in OIM. Together, our data suggest that OIM cells over-activate the process of bone resorption by osteoclasts. In addition, OIM cells showed upregulation of *IL19* (Interleukin 19), which stimulates the differentiation of hematopoietic stem cells into macrophages. Those can further differentiate into osteoclasts under the stimulation of M-CSF (macrophage colony-stimulating factor), and RANK-L.

The production of M-CSF can also be triggered by the presence of defective or misfolded proteins such as homotrimeric type I collagen.^42^ In addition, *CTSD* (Cathepsin D), which is involved in the breakdown of collagen, is upregulated in OIM, further indicating that homotrimeric collagen is being degraded.^43^

Having shown that the presence of the OI-causative mutation negatively impacts the process of differentiation of pre-osteoblasts, a comparative analysis was performed on OIM and WT osteoblasts after three weeks of osteogenic differentiation.

### The TGF-β signalling pathway is upregulated in OIM osteoblasts

Interestingly, the OI mutation does not affect un-differentiated pro-osteoblasts when cultivated in expansion media (**Figure 6**). In contrast, after osteogenic differentiation OIM and WT cell gene expression differs significantly, with 451 genes being downregulated and 455 genes being upregulated in OIM, when compared to WT. These data further confirm that the OI-causative mutation impacts not only pre-osteoblasts differentiation but also modifies the gene expression profile of differentiated osteoblasts. The expression of genes belonging to the TGF-β pathway (such as *TGF-β-R3*) is higher in OIM osteoblasts. Those include *IL6* (Interleukin 6), which contributes to bone resorption through the stimulation of osteoclast activity^44^; *GDF6*, and *CDKN1A* (cyclin-dependent kinase inhibitor 1), which are upregulated in tissue repair after fracture. Simultaneously, we observed that *LTBP4* (latent transforming growth factor beta binding protein 4), which regulates the bioavailability of TGF-β^45^, as well as *BAMBI* (BMP and activin membrane-bound inhibitor), which inhibits signals from the TGF-β superfamily, including bone morphogenetic proteins (BMPs) and activins^46^, were less expressed in OIM osteoblasts.

**Figure 6.**
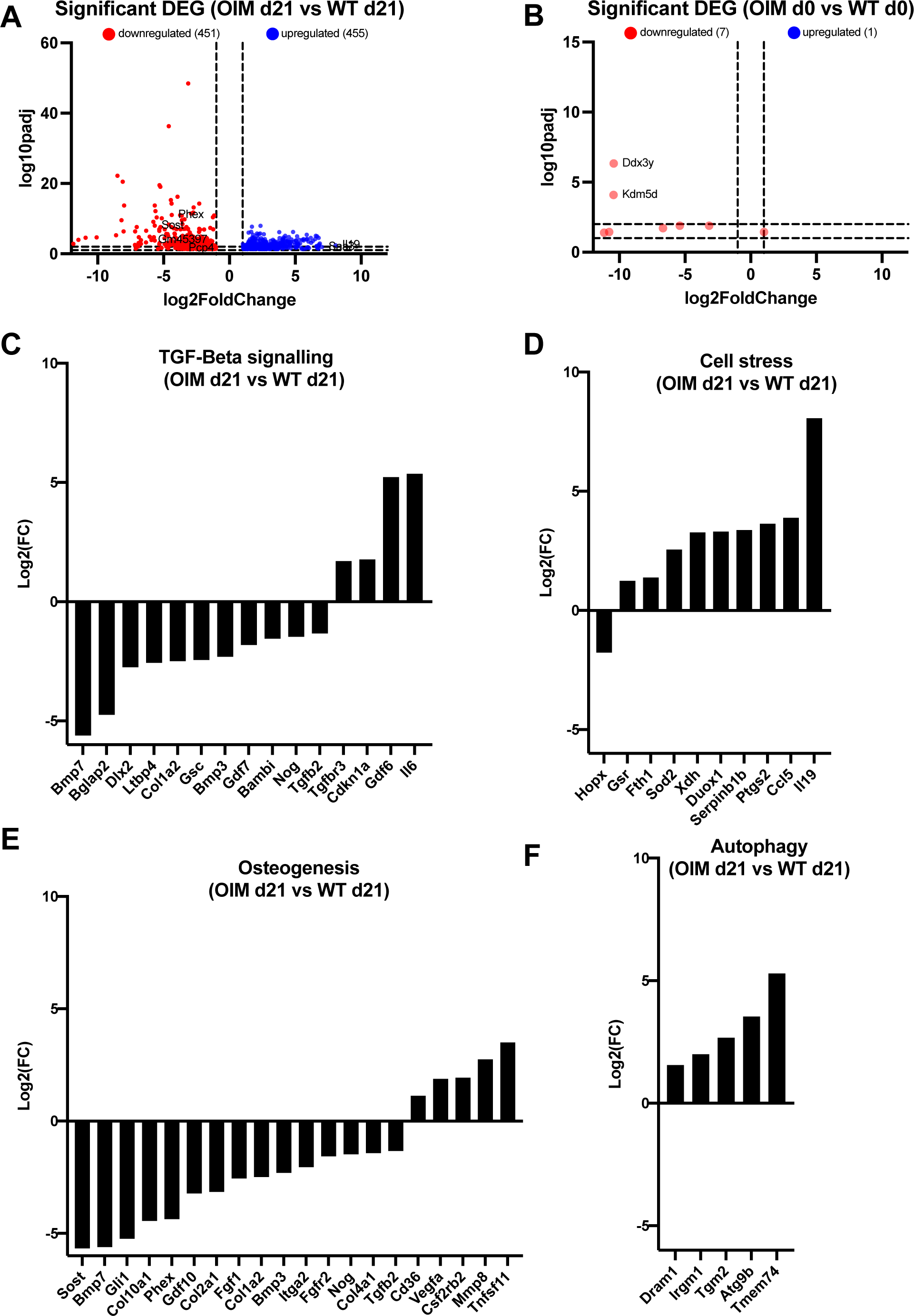
RNAseq analysis of mature WT vs mature OIM osteoblasts and WT pre-osteoblasts vs OIM pre osteoblasts during osteogenic differentiation. (A) Volcano plots showing significant (FDR adjusted p<0.05) DEGs (differentially expressed genes) in mature OIM (OIM d21) vs mature WT (WT d21) (A), and OIM pre-osteoblast (OIM d0) vs WT pre-osteoblasts (WT d0). DEGs were plotted by pathways: TGF-B signalling (C), cell stress (D), osteogenesis (E), autophagy (F).

### OIM osteoblasts do not reach the same level of maturity as WT osteoblasts and remain at an earlier stage of differentiation

This is illustrated by the high expression of genes during the early stage of osteogenic differentiation such as *TNFSF11* (*RANKL*), which is expressed by early osteoblasts and stimulates the differentiation of osteoclasts. In contrast, we observed a downregulation of late osteogenic marker expression such as *BGLAP* and *SOST*, which are expressed by osteocytes^47^ and *GLI1* (glioma associated oncogene homolog 1), which stimulates osteoblast differentiation by promoting the transcription of osteogenesis-related genes.^48^ In addition, we observed a downregulation of *PHEX* (phosphate regulating endopeptidase x-linked), which may also contribute to impair mineralization by altering phosphate levels.^49^

### The phenotype of OIM osteoblasts has a feedback inhibitory effect on osteogenesis and on bone extracellular matrix

*DLX2* (distal-less homeobox 2), a transcription factor which controls the expression of genes involved in osteoblast differentiation^50^; and *BMP7*, which is a potent inducer of osteogenesis, are considerably less expressed in OIM osteoblasts. This indicates that upon differentiation, OIM osteoblasts themselves negatively feedback on osteogenesis, further preventing the progression of pre-osteoblast cells to a more mature differentiation state. Interestingly, *MMP8*, which is involved in the degradation of collagen, is highly expressed in OIM osteoblasts compared to WT cells, possibly due to the presence of defective collagen fibres. *ITGA2* (integrin subunit alpha 2), known to have a specific binding affinity for collagen type I^51^, is down regulated in OIM osteoblasts, which contributes to further compromise bone ECM remodelling by influencing the deposition of minerals and the organization of type I collagen fibres.

### Cell stress signalling pathway is upregulated in OIM osteoblasts

Several genes involved in the cell stress signalling pathway are more highly expressed in OIM osteoblasts than in WT counterparts, likely in response to the abnormal cytoplasmic presence of defective COL1A2 mRNA. For example, these include *IL19* (interleukin 19), which plays an important role in tissue repair and remodelling through its anti-inflammatory effects^52^; *CCL5* (chemokine c-c motif ligand 5), which attracts T cells to the side of inflammation or tissue damage, and *SERPINB1B*, which protects tissues from damage at inflammatory sites.^53^

*DUOX1* (dual oxidase 1) and *XDH* (xanthine dehydrogenase) both play a significant role in the generation of ROS in bones, which can lead to oxidative stress and contributes to negatively impact osteoblast function, including a reduction of bone ECM synthesis and impaired mineralization. Excess ROS also leads to the overactivation of osteoclasts, further unbalancing bone turnover.^54^

### Autophagy is upregulated in OIM osteoblasts

We observed an upregulation of 5 key genes involved in the autophagy signalling pathway, *DRAM1* (DNA damage-regulated autophagy modulator 1), which promotes the sequestration and degradation of cellular components within the autophagosome^55^; *IRGM1* (immunity related GTPase M) and *ATG9B* (autophagy-related protein 9B), which both contribute to the formation of the autophagosome; *TGM2* (transglutaminase 2), expression of which is induced in response to oxidative stress^56^; and *TMEM74* (transmembrane protein 74), an autophagosome protein.^57^

## Discussion

The integrity of the skeletal system is maintained through the constant remodelling of the bone extracellular matrix (ECM) by osteoblasts, through the synthesis of type I collagen and minerals, and by osteoclasts, which are responsible for bone resorption and release of minerals back into the bloodstream. The presence of the recessive OIM-causative variant directly results in the extracellular secretion of abnormal homotrimeric COL1 and intracellularly to the accumulation of COL1A2 mRNA. These two primary effects contribute to abnormal osteoblast development and function, and dysregulated bone remodelling leading to skeletal fragility to homozygous OIM mice. The comparative imaging of 1-week and 8-week-old OIM and WT tibia revealed that by one week of age, several bone parameters are not yet impacted by the recessive OIM-causative variants. This includes (1) the volume of the dense and compact bone (also called cortical bone) which function is to protect the inner trabecular bone and to provide strength and support to the skeleton; (2) the volume of the bone marrow cavity (medullary canal), which influence the bone marrow’s capacity to produce blood cells; (3) the thickness of the cortex, which contributes to the overall strength of bones by providing the ability to withstand mechanical stress and resistance to fractures; (4) the total porosity, which refers to the volume of void space within the bone tissue; (5) the thickness of the trabecular bone, which contributes to the bone’s resistance to fractures, in particular in the regions of high load-bearing, (6) the “intersection surface”, which refers to the surface where trabeculae cross each other to form a three-dimensional lattice-like structure that confers mechanical strength to the trabecular bone by allowing the distribution of forces within the bone, and finally (7) the “trabecular pattern factor”, which gives an overall measure of the bone quality. However, 1-week-old OIM mouse bones already show a decrease in the thickness of the medullary canal, and an increase in the bone volume fraction (BV/TV), bone surface density (BS/TV), trabecular number, and tissue mineral density (TMD). This is possibly due to the visible greater size of the OIM growth plate compared to age-matched WT bones, which is likely a compensatory mechanism in response to the abnormal structure of the collagen matrix. By the age of 8 weeks, skeletal maturity and growth plate closure is almost complete. All parameters except for medullary volume, such as BV/TV, BS/TV and trabecular number demonstrate deterioration by this stage, reflecting an overall picture of reduced trabecular bone.

Together, these results suggests that perinatal interventions could protect the developing OIM bones. The analysis of mineral deposition by OIM and WT osteoblasts showed that the OIM-causative mutation does not impact the quantity of minerals being deposited in the bone matrix, but rather its organization. This is important because mineralization of the collagen network (which provides tensile strength) confers compressive strength to bones and thereby contributes to resistance to fracture. The structure of the homotrimeric COL1 network may directly prevent its proper mineralization, or the incomplete differentiation of OIM pre-osteoblasts may result in the diffuse deposition of minerals in OIM bones.

Whole genome transcriptome analysis revealed that OIM pre-osteoblasts isolated from neonatal calvaria do not differ from their WT counterparts when cultivated in expansion media. However, when exposed to osteogenic permissive conditions over a 3-week period, transcriptome profiling revealed that, OIM cells activate different signalling pathways compared to WT pre-osteoblasts. In addition, OIM and WT mature osteoblasts are phenotypically and genetically different. This is the first study showing how the OIM-causative mutation impacts the whole genome during the process of osteogenic differentiation and compares the transcriptome of OIM and WT pre-osteoblasts and osteoblasts. As previously reported, incomplete osteogenesis was found in OIM cells (**Figure 7**), with 17 genes being exclusively upregulated in WT osteoblasts (*TGFBR2, BMP2, MMP2, SP7, BMP5, FGFR2, COL2A1, BMP3, FDG10, COMP, ITGA2, TNF, SOST, ALP, BGLAP, COL10A1*) and 10 genes being exclusively upregulated in OIM osteoblasts (*TGFB1, FGF2, TGFBR3, MMP9, ITGA3, TNFSF11, MMP8, MMP10, CSF2 and CSF3*), whilst only 2 genes are exclusively downregulated in OIM osteoblasts (*COL5A1* and *SERPINH1*), and one gene in WT osteoblasts (*BMPR1B*). The fact that OIM fail to fully mature but remain at an earlier stage of differentiation, has several consequences, not least the activation of the osteoclast pathway, thus compromising bone remodelling by favouring bone resorption. It also negatively impacts the bone ECM mineralization process (size, distribution, and organization of minerals), thereby contributing to further decreasing bone strength, especially as the structure of the collagen network may already contribute to defective matrix mineralization. In addition, the lower levels of osteocalcin in OIM may have knock-on effects on overall health, since this non-collagenous protein is not only involved in the process of mineralization through its incorporation into the bone matrix and its involvement in the process of hydroxyapatite crystals formation but is also involved in the regulation of calcium homeostasis, glucose metabolism, insulin sensitivity, and metabolism. Our data indicate that the primary intracellular (COL1A2 mRNA accumulation) and extracellular (homotrimeric COL1 lattice) consequences of the homozygous OIM-causative variant trigger a cascade of knock-on effects, both at the level of osteoblast behaviour and matrix structural properties, that together contribute to bone brittleness in a multifactorial manner.

**Figure 7.**
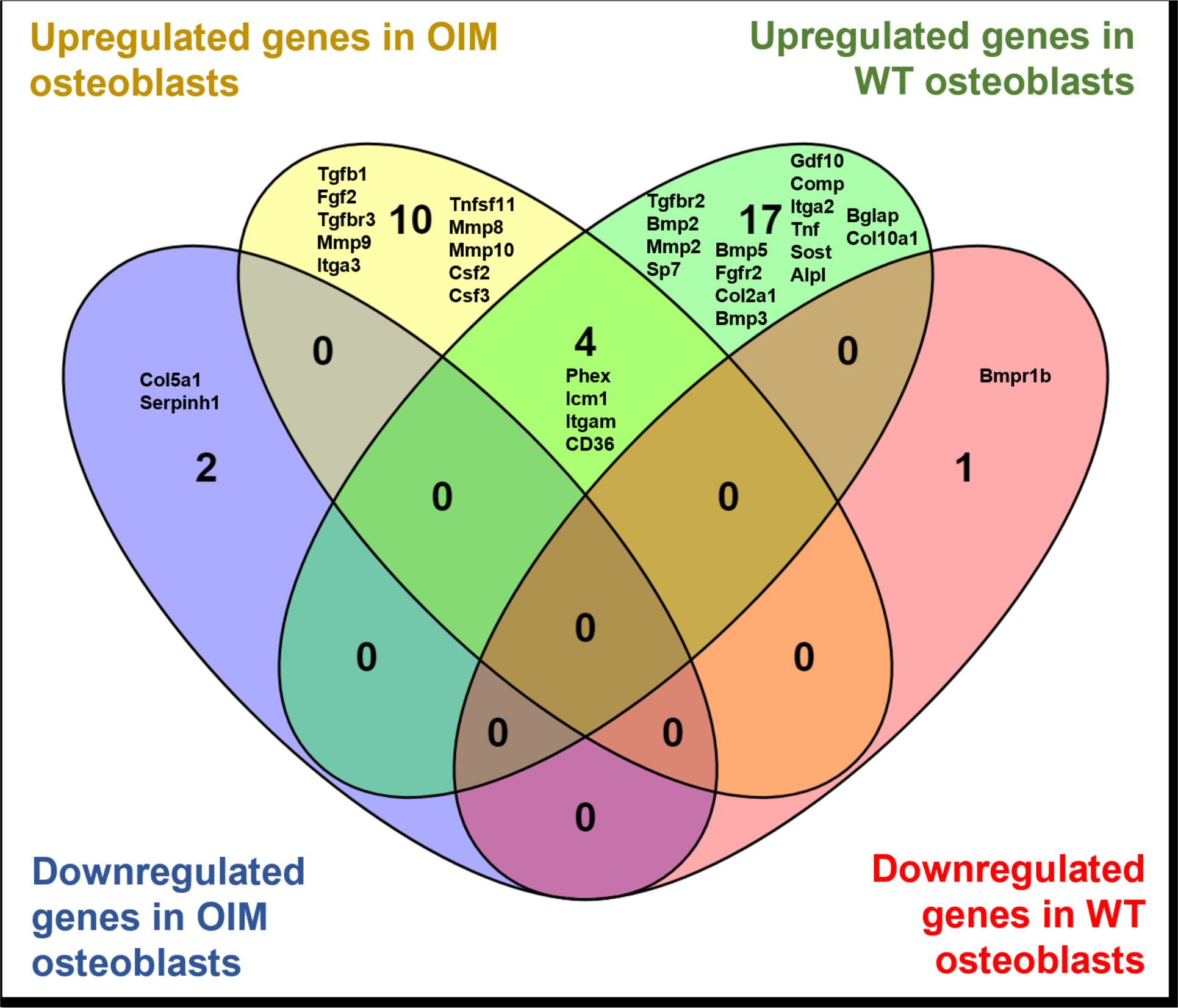
Venn Diagram showing significant DEG in the osteogenic domain co-regulated in WT osteoblasts vs WT pre-osteoblasts (WT d21 vs WT d0) and OIM osteoblasts vs pre-osteoblasts (OIM d21 vs OIM d0)

Figure 8 shows a simplified overview of the several mechanisms that contribute to compromised osteogenesis and result in skeletal fragility. The presence of the structurally defective homotrimeric bone ECM may be directly responsible for the upregulation of the TGF-β pathway. Grafe et al.^58^ previously showed in two different recessive (*Crtap^-/-^*) and dominant (*Col1a2^tm1.1Mcbr^*) mouse models of OI that the reduced binding of the proteoglycan decorin to type I collagen prevented the sequestration of TFG-β in its latent form. The hypothesis of a dysregulated matrix-cell signalling in the OIM model is supported by data showing that homotrimeric collagen has a higher tendency of forming kinks, which leads to larger lateral intermolecular spacing^59^, and that this structural alteration causes reduced intermolecular crosslinking.^60^ Excessive TGF-β signalling can not only stimulate the early stage of osteoblast differentiation, but at the same time prevent the cells from reaching a mature osteoblast phenotype, as well as stimulating inflammation. The latter also has an inhibitory effect on osteogenesis whilst increasing oxidative stress, which is already stimulated by the production of ROS in response to the abnormal intracellular accumulation of *COL1A2* mRNA. Oxidative stress is known to activate autophagy, which also contributes to the inhibition of osteoblast maturation. ROS production also contribute to the degradation of the already structurally compromised bone ECM. The combination of defective collagen matrix and impaired osteoblast maturation leads to defective mineralization of bone ECM, conferring brittleness to the bones. The repetitive occurrence of fractures could contribute to further reactivate the circuitry linking excessive TGF-β signalling, increased inflammation, oxidative stress, and autophagy. Defective osteogenesis and increased bone resorption further weaken the OIM bone ECM.

**Figure 8.**
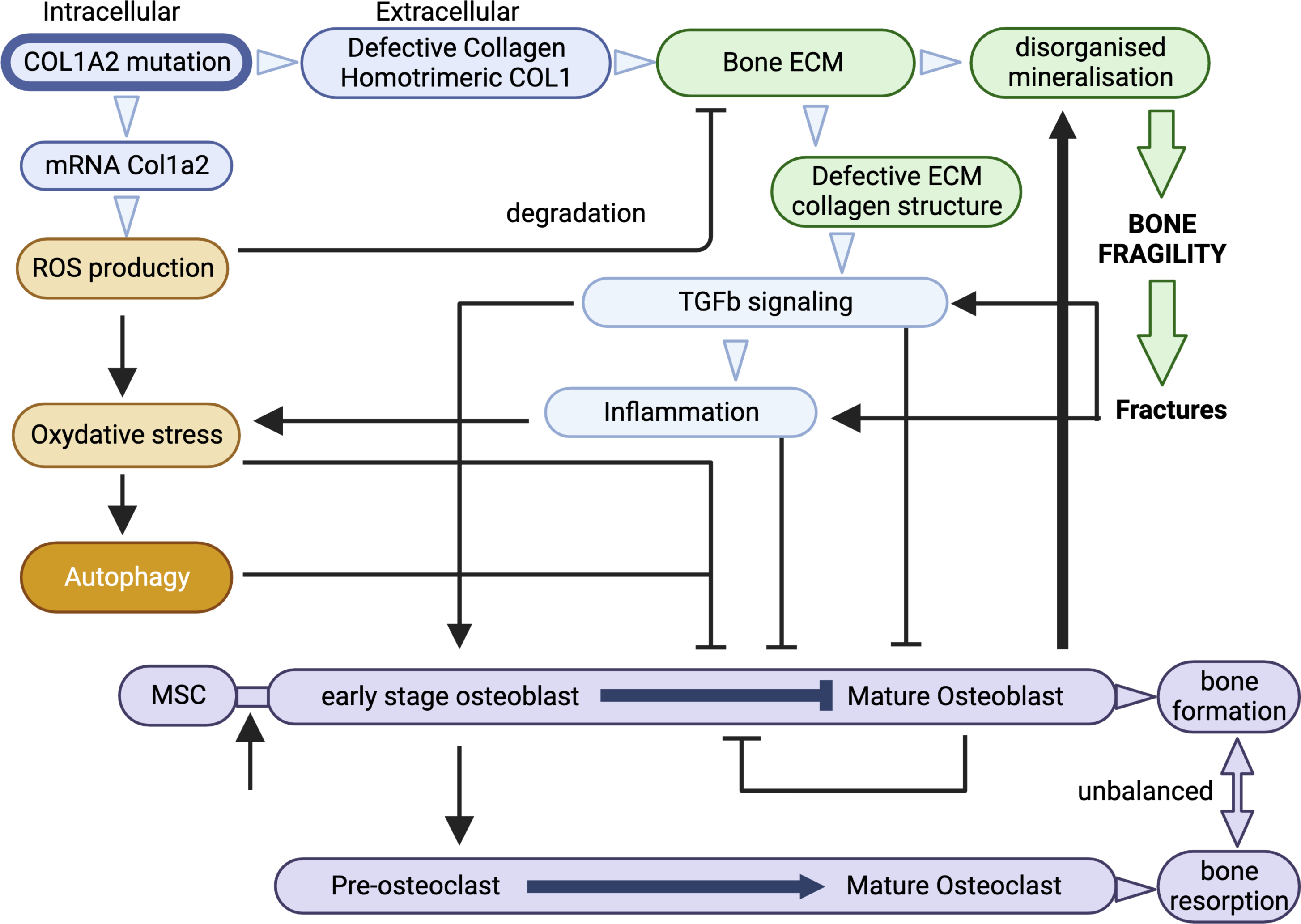
simplified model of the various pathways contributing to bone fragility in homozygous OIM mice. Created by Biorender.

These data provide an overview of the pleiotropic effects in the homozygous OIM mice and give additional insight into the mechanisms of OI disease progression in this model. Our data also identify multiple clinically relevant endpoints to assess the efficacy of innovative therapeutic intervention aiming at improving skeletal health for people with OI.

## Methods

### Animals

All experimental protocols complied with the UK Home Office guidelines (PPL 70/6857). Heterozygous male and female (B6C3Fe a/a-Col1a2oim/Col1a2oim) mice (Jackson Laboratory) were housed in individual ventilated cages at 21□°C with a 12:12-hour light dark cycle. Offspring were genotyped by sequencing the oim fragment then homozygous and wild type colonies established. For the analysis of the neonatal mice, the pups were culled at 7 days postnatal while adult animals were culled at 8 week postnatal.

### Primary osteoblast precursors isolation and expansion

Murine neonatal calvaria were harvested from 4-7 days old OIM and WT mice. The skulls were washed in PBS (Phosphate Buffered Saline) and transferred onto a sterile dish for dissection. A knife blade was used to scrape off any soft tissues. The bone tissue was then sequentially digested in a solution of 0.2% collagenase type II (Gibco) for a total of 1h 30min. The final digest was placed in a plastic cell culture dish with alpha modified Eagle’s medium (alpha-MEM) (Invitrogen) supplemented with 10% fetal bovine serum (Gibco), 2□mM L-glutamine, 50□IU/ml penicillin and 50□mg/ml streptomycin (Gibco), at 37□°C in a 5% CO_2_ incubator. Cells were expanded for about a week until there was a homogeneous population of ALP (alkaline phosphatase) positive cells at subconfluency.

### *In vitro* osteogenic differentiation

Cells were differentiated down the osteoblast lineage for up to 4 weeks in alpha-MEM supplemented with 2□mM β-glycerophosphate, 0.2□mM ascorbic acid and 10^−8^□M dexamethasone, then fixed in 4% paraformaldehyde in PBS.

### Quantification of mineralised bone formation

Cells were stained with a solution of 1% Alizarin Red to visualize the calcium and phosphate deposits. Once fully dry, the plates were then scanned on a high-resolution flat-bed scanner (Epson Perfection v600). ImageJ free software was used to measure the area fraction (%) of the mineralised bone nodules.

### Micro-computed tomographic analysis

Tibiae were isolated from 1-week-old homozygous oim mice (n=7), 1-week-old wild type (WT) mice (n=7), 8-week-old homozygous OIM mice (n□=□6) and 8-weeks old wild type (WT) mice (n□=□6). The bones were fixed in 10% neutral buffered formalin for 24□h, then washed in phosphate buffered saline and stored in 70% ethanol until scanning. All the scans were performed using a Skyscan 1172□μ-CT scanner (Bruker, Belgium). The bones were scanned at 50□kV and 200□μA using a 0.5□mm aluminium filter and a pixel resolution of 4.3□μm (for the 8-weeks old mice) and 3.3 μm (for the 7-days old pups). To analyse the trabecular bone, a region of interest of the length of 0.8□mm was selected below the growth plate of the proximal tibia. To analyse cortical bone, a 0.4□mm long region of interest was selected with an offset of 2□mm below the growth plate. The images were reconstructed using the Skyscan NRecon software and the following cortical and trabecular morphometric parameters were calculated using the Skyscan CT Analyzer (CTAn) software: percent bone volume (BV/TV) (%), bone surface density (BS/TV), trabecular thickness (mm), trabecular number (mm^−1^), trabecular pattern factor (mm^−1^), intersection surface (mm^2^), cortical volume (mm^3^), medullary volume (mm^3^), cortical thickness (mm), total porosity (%), medullary canal thickness (mm), and tissue mineral density (TMD) (g/cm^3^). Bone mineral density was measured using 0.25 and 0.75□g/cm^3^ calcium hydroxyapatite calibration phantoms (Bruker) as a reference. P values were calculated using analysis of variance (one-way ANOVA) followed by Bonferroni’s multiple comparison post hoc test. Differences with a P-value of < 0.05 were considered significant. Data were expressed as mean□±□SEM (standard error of the mean).

### Transcriptomic analysis (RNA-seq)

Total RNA was extracted from oim and wt pre-osteoblasts and mature osteoblasts using TRIzol (Invitrogen). RNA sequencing (RNA-seq) analysis was performed (Genewiz), to investigate gene-expression profiling in the following groups (OIM pre-osteoblast, WT pre-osteoblast, OIM osteoblast, WT osteoblast). Differential gene expression analyses for all the group comparisons were performed using DESeq2 open-source software. The Wald test was used to generate p-values and log2 fold changes. Genes with a p-adjusted value < 0.05 and absolute log2 fold change > 1 were called as differentially expressed genes for each comparison.

## Author contributions

Conceptualisation and methodology, M.C. and P.V.G.; Validation, M.C. and P.V.G.; Formal analysis, M.C. and P.V.G.; Investigation, M.C., R.S., E.P., A.L.D., R.S., P.V.G.; Provision of material, P.V.G.; Writing, M.C. and P.V.G.; Review, Editing & Acceptance of final manuscript, all authors; Visualisation, P.V.G.; Supervision, P.V.G.; Funding Acquisition, P.V.G.

## Acknowledgements.

This study was supported by grants from Medical Research Council, Rosetrees Trust and The Stonygate Trust.

## References

1. Le, B. Q. et al. The Components of Bone and What They Can Teach Us about Regeneration. Materials 11, 14 (2017).

2. Dijk, F. S. V. & Sillence, D. O. Osteogenesis imperfecta: Clinical diagnosis, nomenclature and severity assessment. Am. J. Méd. Genet. Part A 164, 1470–1481 (2014).

3. Chipman, S. D. et al. Defective pro alpha 2(I) collagen synthesis in a recessive mutation in mice: a model of human osteogenesis imperfecta. Proc Natl Acad Sci U S A 90, 1701–1705 (1993).

4. Alcorta-Sevillano, N., Infante, A., Macías, I. & Rodríguez, C. I. Murine Animal Models in Osteogenesis Imperfecta: The Quest for Improving the Quality of Life. Int. J. Mol. Sci. 24, 184 (2022).

5. Li, H. et al. Immature osteoblast lineage cells increase osteoclastogenesis in osteogenesis imperfecta murine. Am J Pathol 176, 2405–2413 (2010).

6. Kalajzic, I. et al. Osteoblastic response to the defective matrix in the osteogenesis imperfecta murine (oim) mouse. Endocrinology 143, 1594–1601 (2002).

7. Grafe, I. et al. Excessive transforming growth factor-beta signaling is a common mechanism in osteogenesis imperfecta. Nat Med 20, 670–675 (2014).

8. Song, I.-W. et al. Targeting transforming growth factor-β (TGF-β) for treatment of osteogenesis imperfecta. J. Clin. Investig. 132, e152571 (2022).

9. Mirigian, L. S. et al. Osteoblast Malfunction Caused by Cell Stress Response to Procollagen Misfolding in alpha2(I)-G610C Mouse Model of Osteogenesis Imperfecta. J Bone Miner Res 31, 1608–1616 (2016).

10. Chipman, S. D. et al. Defective pro alpha 2(I) collagen synthesis in a recessive mutation in mice: a model of human osteogenesis imperfecta. Proc Natl Acad Sci U S A 90, 1701–1705 (1993).

11. Guillot, P. V. et al. Intrauterine transplantation of human fetal mesenchymal stem cells from first-trimester blood repairs bone and reduces fractures in osteogenesis imperfecta mice. Blood 111, 1717–1725 (2008).

12. Li, J., Yu, L., Guo, S. & Zhao, Y. Identification of the molecular mechanism and diagnostic biomarkers in the thoracic ossification of the ligamentum flavum using metabolomics and transcriptomics. BMC Mol. Cell Biol. 21, 37 (2020).

13. Takata, T., Araki, S., Tsuchiya, Y. & Watanabe, Y. Oxidative Stress Orchestrates MAPK and Nitric-Oxide Synthase Signal. Int. J. Mol. Sci. 21, 8750 (2020).

14. Indovina, P., Pentimalli, F., Casini, N., Vocca, I. & Giordano, A. RB1 dual role in proliferation and apoptosis: Cell fate control and implications for cancer therapy. Oncotarget 6, 17873–17890 (2015).

15. Crighton, D., Wilkinson, S. & Ryan, K. M. DRAM Links Autophagy to p53 and Programmed Cell Death. Autophagy 3, 72–74 (2007).

16. Wang, S. et al. Upregulation of ATG9b by propranolol promotes autophagic cell death of hepatic stellate cells to improve liver fibrosis. J. Cell. Mol. Med. (2023) doi:10.1111/jcmm.18047.

17. Zeng, L. et al. SDC1-TGM2-FLOT1-BHMT complex determines radiosensitivity of glioblastoma by influencing the fusion of autophagosomes with lysosomes. Theranostics 13, 3725–3743 (2023).

18. Mo, H. et al. C-X-C Chemokine Receptor Type 4 Plays a Crucial Role in Mediating Oxidative Stress-Induced Podocyte Injury. Antioxid. Redox Signal. 27, 345–362 (2017).

19. Sun, Y. et al. TMEM74 promotes tumor cell survival by inducing autophagy via interactions with ATG16L1 and ATG9A. Cell Death Dis. 8, e3031–e3031 (2017).

20. Ho, H., Cheng, M. & Chiu, D. T. Glucose-6-phosphate dehydrogenase – from oxidative stress to cellular functions and degenerative diseases. Redox Rep. 12, 109–118 (2007).

21. Muñoz-Sánchez, J. & Chánez-Cárdenas, M. E. A Review on Hemeoxygenase-2: Focus on Cellular Protection and Oxygen Response. Oxidative Med. Cell. Longev. 2014, 604981 (2014).

22. Wang, Y., Branicky, R., Noë, A. & Hekimi, S. Superoxide dismutases: Dual roles in controlling ROS damage and regulating ROS signaling. J. Cell Biol. 217, 1915– 1928 (2018).

23. Redza-Dutordoir, M. & Averill-Bates, D. A. Interactions between reactive oxygen species and autophagy Special issue: Death mechanisms in cellular homeostasis. Biochim. Biophys. Acta (BBA) - Mol. Cell Res. 1868, 119041 (2021).

24. Kim, Y. C. & Guan, K.-L. mTOR: a pharmacologic target for autophagy regulation. J. Clin. Investig. 125, 25–32 (2015).

25. Song, I.-W. et al. Targeting transforming growth factor-β (TGF-β) for treatment of osteogenesis imperfecta. J. Clin. Investig. 132, e152571 (2022).

26. Wu, M., Chen, G. & Li, Y.-P. TGF-β and BMP signaling in osteoblast, skeletal development, and bone formation, homeostasis and disease. Bone Res. 4, 16009 (2016).

27. Caplan, A. I. & Correa, D. PDGF in bone formation and regeneration: New insights into a novel mechanism involving MSCs. J. Orthop. Res. 29, 1795–1803 (2011).

28. Kenner, L. et al. Mice lacking JunB are osteopenic due to cell-autonomous osteoblast and osteoclast defects. J. Cell Biol. 164, 613–623 (2004).

29. Shen, B. et al. BMP-13 Emerges as a Potential Inhibitor of Bone Formation. Int. J. Biol. Sci. 5, 192–200 (2009).

30. Zhou, Y. et al. The miR-204-5p/FOXC1/GDF7 axis regulates the osteogenic differentiation of human adipose-derived stem cells via the AKT and p38 signalling pathways. Stem Cell Res. Ther. 12, 64 (2021).

31. Lewiecki, E. M. Role of sclerostin in bone and cartilage and its potential as a therapeutic target in bone diseases. Ther. Adv. Musculoskelet. Dis. 6, 48–57 (2014).

32. Felber, K., Elks, P. M., Lecca, M. & Roehl, H. H. Expression of osterix Is Regulated by FGF and Wnt/β-Catenin Signalling during Osteoblast Differentiation. PLoS ONE 10, e0144982 (2015).

33. Widmer, C. et al. Molecular basis for the action of the collagen-specific chaperone Hsp47/SERPINH1 and its structure-specific client recognition. Proc. Natl. Acad. Sci. 109, 13243–13247 (2012).

34. Drezner, M. K. PHEX gene and hypophosphatemia. Kidney Int. 57, 9–18 (2000).

35. Docheva, D., Popov, C., Alberton, P. & Aszodi, A. Integrin signaling in skeletal development and function. Birth Defects Res. Part C: Embryo Today: Rev. 102, 13– 36 (2014).

36. Troilo, H. et al. The role of chordin fragments generated by partial tolloid cleavage in regulating BMP activity. Biochem. Soc. Trans. 43, 795–800 (2015).

37. Katagiri, T. & Watabe, T. Bone Morphogenetic Proteins. Cold Spring Harb. Perspect. Biol. 8, a021899 (2016).

38. Li, X., Jin, L. & Tan, Y. Different roles of matrix metalloproteinase 2 in osteolysis of skeletal dysplasia and bone metastasis. Mol. Med. Rep. 23, 70 (2021).

39. Luchian, I., Goriuc, A., Sandu, D. & Covasa, M. The Role of Matrix Metalloproteinases (MMP-8, MMP-9, MMP-13) in Periodontal and Peri-Implant Pathological Processes. Int. J. Mol. Sci. 23, 1806 (2022).

40. Zhang, H. et al. MMP9 protects against LPS-induced inflammation in osteoblasts. Innate Immun. 26, 259–269 (2019).

41. Kohli, S. S. & Kohli, V. S. Role of RANKL–RANK/osteoprotegerin molecular complex in bone remodeling and its immunopathologic implications. Indian J. Endocrinol. Metab. 15, 175–181 (2011).

42. Tojo, N., Asakura, E., Koyama, M., Tanabe, T. & Nakamura, N. Effects of macrophage colony-stimulating factor (M-CSF) on protease production from monocyte, macrophage and foam cell in vitro: a possible mechanism for anti-atherosclerotic effect of M-CSF. Biochim. Biophys. Acta (BBA) - Mol. Cell Res. 1452, 275–284 (1999).

43. Fox, C. et al. Inhibition of lysosomal protease cathepsin D reduces renal fibrosis in murine chronic kidney disease. Sci. Rep. 6, 20101 (2016).

44. Smith, J. K. IL-6 and the dysregulation of immune, bone, muscle, and metabolic homeostasis during spaceflight. npj Microgravity 4, 24 (2018).

45. Rifkin, D. et al. The role of LTBPs in TGF beta signaling. Dev. Dyn. 251, 75–84 (2022).

46. Fan, Y. et al. BAMBI Elimination Enhances Alternative TGF-β Signaling and Glomerular Dysfunction in Diabetic Mice. Diabetes 64, 2220–2233 (2015).

47. Wang, J. S., Mazur, C. M. & Wein, M. N. Sclerostin and Osteocalcin: Candidate Bone-Produced Hormones. Front. Endocrinol. 12, 584147 (2021).

48. Shi, Y. et al. Gli1 identifies osteogenic progenitors for bone formation and fracture repair. Nat. Commun. 8, 2043 (2017).

49. Martin, A. et al. Bone proteins PHEX and DMP1 regulate fibroblastic growth factor Fgf23 expression in osteocytes through a common pathway involving FGF receptor (FGFR) signaling. FASEB J 25, 2551–62 (2011).

50. Zhang, J., Zhang, W., Dai, J., Wang, X. & Shen, S. G. Overexpression of Dlx2 enhances osteogenic differentiation of BMSCs and MC3T3-E1 cells via direct upregulation of Osteocalcin and Alp. Int. J. Oral Sci. 11, 12 (2019).

51. Rattanasinchai, C., Navasumrit, P. & Ruchirawat, M. Elevated ITGA2 expression promotes collagen type I-induced clonogenic growth of intrahepatic cholangiocarcinoma. Sci. Rep. 12, 22429 (2022).

52. Jain, S., Gabunia, K., Kelemen, S. E., Panetti, T. S. & Autieri, M. V. The Anti-Inflammatory Cytokine Interleukin 19 Is Expressed By and Angiogenic for Human Endothelial Cells. Arter., Thromb., Vasc. Biol. 31, 167–175 (2011).

53. Kassem, D. H., Adel, A., Sayed, G. H. & Kamal, M. M. A Novel SERPINB1 Single-Nucleotide Polymorphism Associated With Glycemic Control and β-Cell Function in Egyptian Type 2 Diabetic Patients. Front. Endocrinol. 11, 450 (2020).

54. Agidigbi, T. S. & Kim, C. Reactive Oxygen Species in Osteoclast Differentiation and Possible Pharmaceutical Targets of ROS-Mediated Osteoclast Diseases. Int. J. Mol. Sci. 20, 3576 (2019).

55. Zhang, R. et al. Deficiency in the autophagy modulator Dram1 exacerbates pyroptotic cell death of Mycobacteria-infected macrophages. Cell Death Dis. 11, 277 (2020).

56. Li, X. et al. Thiol oxidative stress-dependent degradation of transglutaminase2 via protein S-glutathionylation sensitizes 5-fluorouracil therapy in 5-fluorouracil-resistant colorectal cancer cells. Drug Resist. Updat. 67, 100930 (2023).

57. Yu, C. et al. TMEM74, a lysosome and autophagosome protein, regulates autophagy. Biochem Biophys Res Commun 369, 622–9 (2008).

58. Grafe, I. et al. Excessive transforming growth factor-beta signaling is a common mechanism in osteogenesis imperfecta. Nat Med 20, 670–675 (2014).

59. Chang, S.-W., Shefelbine, S. J. & Buehler, M. J. Structural and Mechanical Differences between Collagen Homo- and Heterotrimers: Relevance for the Molecular Origin of Brittle Bone Disease. Biophys. J. 102, 640–648 (2012).

60. Fang, M. & Holl, M. M. B. Variation in type I collagen fibril nanomorphology: the significance and origin. BoneKEy Rep. 2, 394 (2013).

